# A Multi-Layered Transcriptomic Analysis Sheds Light on Antisense Transcription, RNA Processing, and SmAP Interactions in *Sulfolobus acidocaldarius*

**DOI:** 10.1101/2025.07.28.667248

**Authors:** Michel Brück, Michael Daume, Lennart Randau, José Vicente Gomes-Filho

## Abstract

The archaeal domain contains organisms that are well-adapted to extreme conditions and changes in their habitat. Post-transcriptional regulation plays a key role in environmental adaptation, including rapid molecular responses to stress conditions. To understand the importance of RNA-based post-transcriptional regulation for these processes, a comprehensive analysis of the presence and processing of regulatory RNAs, as well as their interactions with other RNAs and proteins, is indispensable. Here, we combine the analysis of several RNA sequencing approaches to reveal the presence of a set of novel non-coding RNAs (ncRNAs), their expression in various conditions, processing, and molecular interactions in the transcriptome of *Sulfolobus acidocaldarius*, a model organism for Archaea. We expand its annotation by 160 intergenic ncRNAs (sRNAs) and 990 antisense RNAs (asRNAs), add the location and motifs of over 6000 transcript processing sites, and determine the interaction of transcripts with Sm-like archaeal proteins (SmAPs). We determined the correlation between the expression patterns of asRNAs and their cognate mRNAs, suggesting transcript-based regulation patterns in gene expression, particularly in response to changing environmental conditions. Additionally, we observed differential binding preferences of SmAP1 and SmAP2 towards mRNA and ncRNAs, suggesting a distribution of regulating roles of these chaperones. Finally, we provide an overview of our post-transcriptional data analysis results, optimized for custom exploration, in the form of a web-based transcriptome atlas (https://vicentebr.github.io/Sulfolobus_atlas/).

**IMPORTANCE:** Post-transcriptional regulation is a key control layer in gene expression. Yet, resources integrating asRNAs, RNA processing sites, and RNA-protein interactions are scarce for archaeal organisms. Here we combine multiple RNA-seq strategies and RIP-seq to expand the *Sulfolobus acidocaldarius* transcriptome with 990 asRNAs, thousands of transcript processing sites, and the interactomes of the essential RNA chaperones SmAP1 and SmAP2. Integrating the novel generated data for the re-analysis of heat-shock transcriptomics and proteomics reveals locus-specific antisense mechanisms that might help to explain the weak correlation between cellular RNA and protein levels reported previously. Moreover, our publicly accessible web atlas provides a community platform to explore these datasets and assist in the formulation of new hypotheses about archaeal RNA regulation.

## INTRODUCTION

Archaea, commonly regarded as the third domain of life, exhibit remarkable adaptability to extreme environments. *Sulfolobus acidocaldarius* is a model organism that optimally grows at 75 °C and pH levels between 2 and 3 (1, 2). Since its first isolation, this organism has been used for a wide variety of genetic and functional studies (3–6). The relative ease of growth and genetic manipulation makes it a model organism for studying archaea and adaptations to extreme environments (7). Despite this, the annotation and functional characterization of ncRNAs and small protein-coding genes in *S. acidocaldarius* remains limited. Multiple studies have highlighted the complexity of archaeal transcriptomes, revealing numerous ncRNAs involved in various regulatory processes, such as transposition, nitrogen metabolism, and oxidative stress response (8–10).

asRNAs, a subset of transcripts that are transcribed on the opposite strand of a given gene, act as important regulators in both prokaryotes and eukaryotes (11–13). For instance, in *Saccharolobus solfataricus*, transposon-derived asRNAs have been shown to regulate phosphate transporter genes, highlighting the potential for asRNA-mediated control in archaeal systems (14). Moreover, in bacteria, antisense transcription is proposed to drive genome-wide mRNA processing by RNase III recruitment (15). In *S. acidocaldarius*, the lack of an RNase III homologue potentially leads to the accumulation of an RNA duplex that is involved in biofilm regulation (16). Although lacking an apparent machinery for RNA duplex degradation, *S. acidocaldarius* contains diverse RNases that are involved in the maturation and degradation of multiple transcripts (17–19).

Post-transcriptional regulation in archaea is further influenced by the presence of RNA chaperones such as Sm-like archaeal proteins (SmAPs), L7Ae, and TRAM (5, 20–23). These proteins are integral to RNA metabolism, participating in processes such as RNA splicing, degradation, and ribonucleoprotein complex assembly (24).

In *S. acidocaldarius*, three SmAP homologs have been identified: SmAP1, SmAP2, and SmAP3. SmAPs typically assemble into homo-oligomeric rings, forming heptameric structures that bind RNA molecules through a conserved pocket selective for uridine-rich sequences (20–22, 25–27). While the exact extent of RNA interaction partners and the functional outcome of such interactions is unknown in *S. acidocaldarius*, studies in related species suggest that these proteins interact with various components of the RNA processing machinery. For example, in *Sa. solfataricus*, SmAP1, and SmAP2 co-purify with proteins involved in RNA processing and degradation, including components of the exosome (27). Thus, this association suggests a role for SmAPs in regulating RNA stability and turnover.

Additionally, structural analyses of SmAPs from *S. acidocaldarius* have provided insights into their RNA-binding properties. Crystallographic studies reveal that these proteins share a common fold with eukaryotic Lsm proteins, forming rings with a central pore that accommodates the RNA. The conservation of this architecture across domains highlights an important role of SmAPs in RNA metabolism (28).

Studies in other archaea, such as *Halobacterium salinarum*, have demonstrated the extensive role of post-transcriptional regulation in environmental adaptation. In *H. salinarum*, a genome-scale atlas revealed that 54% of protein-coding genes are targeted by multiple post-transcriptional mechanisms, including SmAP1 binding, asRNAs, and RNase-mediated processing, reinforcing a highly complex regulatory scenario (29). Thus, understanding the interplay between asRNAs and SmAPs, as well as the patterns of RNA degradation, is crucial for elucidating the post-transcriptional regulatory networks in *S. acidocaldarius*.

By combining (i) the detection and annotation of novel ncRNAs, including asRNAs; (ii) differential expression analyses of ncRNAs and mRNAs; (iii) transcriptome wide mapping of transcript processing sites (TPSs); and (iv) the characterization of binding partners of SmAP1 and SmAP2, this study provides a comprehensive view of the *S. acidocaldarius* DSM 639 transcriptome. Moreover, the resulting datasets are consolidated into an interactive web-based platform that facilitates data exploration and hypothesis generation.

## RESULTS

### Comprehensive expansion of transcript annotation in *S. acidocaldarius*

The complete genome of *S. acidocaldarius* DSM 639 (hereafter referred to as *S. acidocaldarius*) was initially published in 2005, revealing a single circular chromosome of approximately 2.23 Mbp (30). As of March 29, 2024, the RefSeq annotation provides a total of 2,373 genes, of which 2,293 are protein-coding and 80 are non-coding (tRNAs, rRNAs, RNase P, pseudogenes). This indicates a lack of annotated regulatory ncRNAs for this species. In prokaryotic genomes, the annotation of ncRNAs, such as intergenic ncRNAs and asRNAs, as well as small protein-coding genes encoding proteins shorter than 50 amino acids, remains a relevant challenge (31, 32).

To address this, we developed an integrated methodology that utilizes previously published transcription start sites (TSS), identified by differential RNA-sequencing (dRNA-seq), and a genome-wide map of RNA termini generated by Term-seq (3, 33). By applying this strategy, we were able to detect and annotate 990 asRNAs, 160 sRNAs, and 95 putative small protein-coding genes. Moreover, to create the final annotation, we curated previously published data (5, 16) to incorporate known small nucleolar RNAs (snoRNAs) and sRNAs, and to remove duplicated entries (File S1). By examining the upstream regions of the novel annotated asRNAs and sRNAs, a common promoter sequence characterized by a TATA motif located 24 to 30 base pairs upstream of the TSS was identified (Fig. 1A), suggesting the determination of primary transcripts.

**Figure 1:**
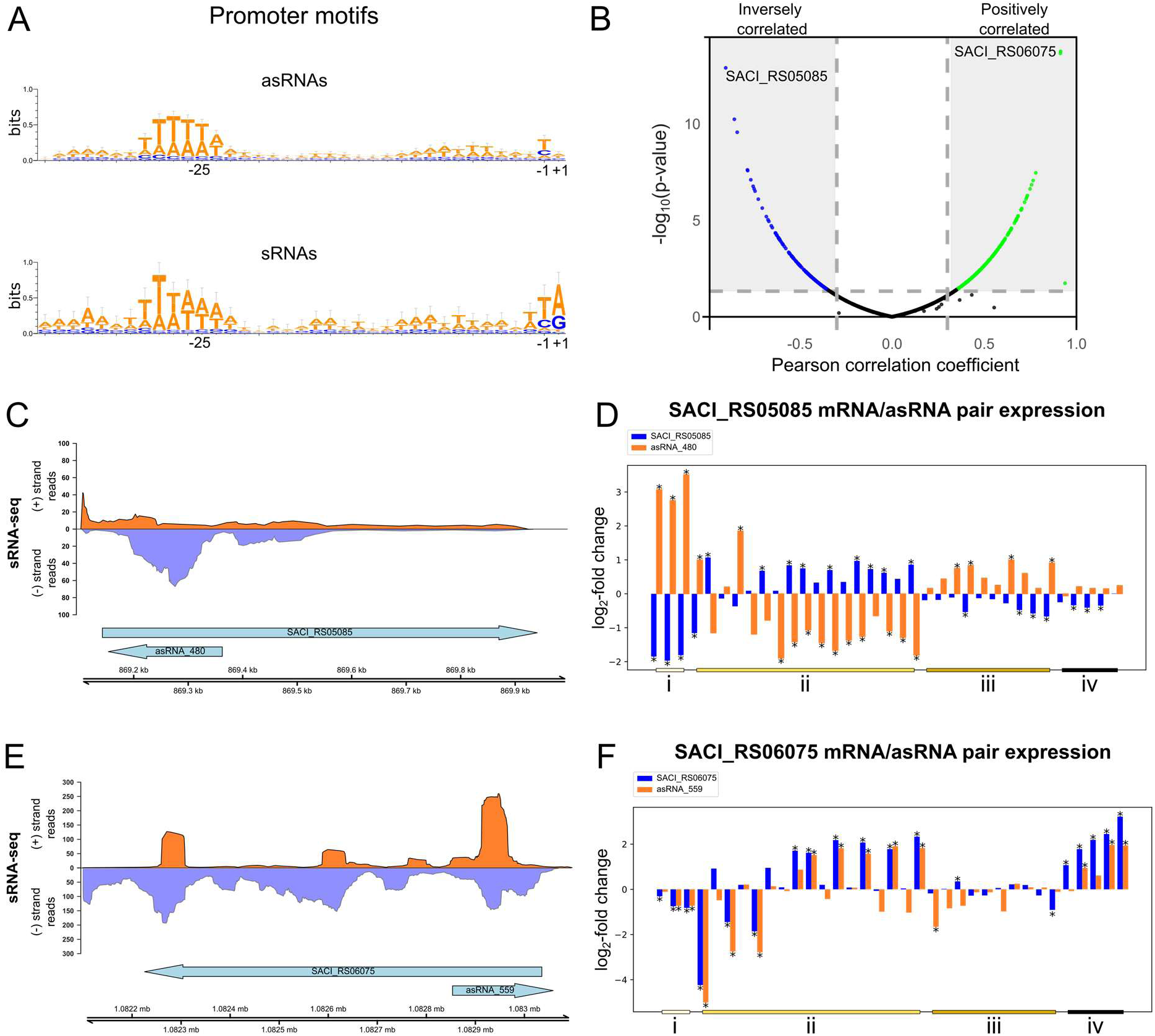
Identification and analysis of novel ncRNAs in *S. acidocaldarius.* (A) Sequence logo of conserved promoter motifs from identified asRNAs and sRNAs, generated using WebLogo3(79). (B) Correlation analysis of asRNAs and their cognate mRNAs. Pairs with a Pearson correlation coefficient (PCC) ≥ 0.5 (positive correlation) or ≤ −0.5 (negative correlation) are shown in green and blue, respectively; uncorrelated or weakly correlated pairs are shown in black. (C) Normalized sRNA-seq coverage plot showing read accumulation at the asRNA_559 locus (− strand). Coverage for the (+) and (−) strands is shown in dark orange and light purple, respectively. Arrows indicate gene orientation: right-facing for (+) strand, left-facing for (−) strand. (D) Differential expression profiles of the positively correlated SACI_RS06075/asRNA_559 pair (PCC = 0.91). (E) Normalized sRNA-seq coverage plot showing read accumulation at the asRNA_480 locus (+ strand), with strand coverage and gene orientation as in panel D. (F) Differential expression and sRNA-seq coverage of the negatively correlated SACI_RS05085/asRNA_480 pair (PCC = −0.9). Experimental conditions include: (i) post-heat shock (86 °C), (ii) varying growth phases, temperature, and pH, (iii) NUDIX gene knockouts, and (iv) post-starvation. Statistically significant changes (p-value < 0.05) are marked with an asterisk (*).

### Differential expression analysis reveals the underlying correlation dynamics between asRNAs and their cognate mRNAs

To provide hints about the functional roles of the identified ncRNAs, we gathered publicly available transcriptomic datasets from *S. acidocaldarius* generated in diverse genetic and environmental contexts: (i) temperature, pH, and growth phase variation; (ii) NUDIX proteins knockout (34); (iii) Post heat stress (86 °C) (35); (iv) Post nutrient limitation (36); and (v) sRNA-seq (5) (Table S1). After data collection, all samples were quality checked, trimmed, and aligned to the genome of *S. acidocaldarius* (see Methods). To calculate differential expression, our novel annotation was used in combination with featureCounts (37) and DESeq2 (38) (File S2). Next, ncRNAs that had log_2_-fold change ≥ 1 or ≤ −1 and p-value < 0.05 were qualified as differentially expressed. Since asRNAs are important players in post-transcriptional regulation in archaea (11, 13, 14, 16, 39), we dedicated a deeper focus on their properties, interactions, partners, and patterns of expression. In total, 828 genes were found to have between one and five associated asRNAs (Table S2).

Utilizing the previously described expression datasets (File S2), we calculated the Pearson Correlation Coefficient (PCC) for each asRNA/mRNA pair. Some mRNA/asRNA pairs exhibited positively correlated expression patterns (PCC ≥ 0.5), meaning the simultaneous increase or decrease in their RNA abundance, whereas another subset displayed inversely correlated expression patterns (PCC ≤ −0.5) (Fig. 1B). For instance, asRNA_480 (Fig. 1C) exhibited significant inverse correlation (PCC = −0.9), being notably up- or downregulated while its cognate mRNA (SACI_RS05085; predicted membrane protein/putative permease of an uncharacterized ABC transporter) was significantly up- or downregulated across the examined conditions (Fig. 1D). Conversely, asRNA_559 (Fig. 1E) demonstrated a consistently significant positive correlation (PCC = 0.91) with its cis-transcribed mRNA (SACI_RS06075; SulA sulfolobicin component (40)) under most tested conditions (Fig. 1F). These findings highlight the intricate interplay between antisense RNAs and their corresponding mRNAs, suggesting that this interaction may influence other post-transcriptional regulation processes (11, 14, 15, 24).

### Identification and mapping of transcriptome-wide processing sites using differential RNA-seq

Emerging evidence indicates that asRNAs, alongside other post-transcriptional mechanisms, may contribute to the maturation and degradation of a wide range of transcripts in archaea (8, 39, 41, 42). To investigate this layer of regulation in *S. acidocaldarius*, we reanalyzed dRNA-seq data to perform a transcriptome-wide identification of TPSs (3, 41). To perform TPS identification, we utilized the Transcription Start Site Annotation Regime Web Service (TSSAR), a statistical framework that identifies significantly depleted 5′ ends in TEX-treated libraries, as previously established for archaeal transcriptome studies (43). Briefly, a high number of reads starting in a specific position in the -TEX samples, which are significantly depleted in +TEX libraries, suggests the presence of 5′ monophosphorylated transcripts (Fig. 2A), a common outcome of RNA processing and degradation events in prokaryotes (44, 45). This analysis identified a total of 6,013 putative TPSs across the *S. acidocaldarius* transcriptome (File S3).

**Figure 2:**
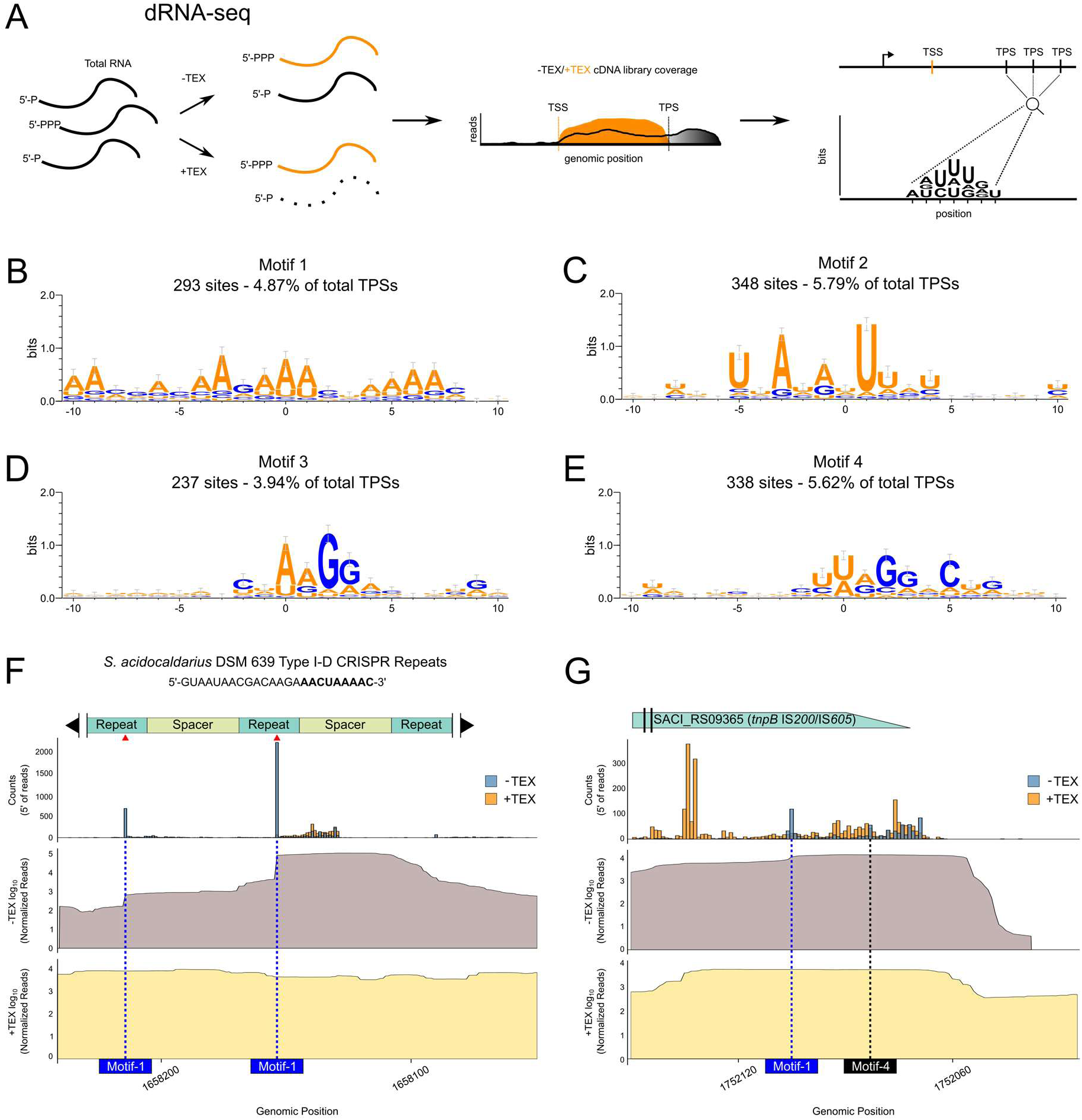
Transcriptome-wide identification of TPSs using dRNA-seq. (A) Schematic overview of the dRNA-seq approach based on Terminator 5′-phosphate-dependent exonuclease (TEX) treatment. The comparison of TEX-treated (+TEX - black coverage) and untreated (−TEX - orange coverage) libraries allows detection of TPSs, using an inverted input strategy relative to transcription start site (TSS) identification. (B-E) Sequence logos of four significantly enriched TPS-associated motifs identified by MEME and generated with WebLogo3 (79). Position 0 indicates the predicted processing site. (F) Mapping of 5′ ends at the *S. acidocaldarius* CRISPR Type I-D array reveals multiple TPSs overlapping conserved repeat sequences. The expected repeat tag is marked in bold in the repeat sequence. Bar plots show normalized 5′ end read counts from +TEX/−TEX libraries. Additionally, coverage plots of log_10_(Normalized Reads) from -TEX (dark gray) and +TEX (light yellow) show peak depletion at the positions where TPSs are identified. Processing events that coincide with Motif-1 sites and match Cas6 cleavage positions are indicated by red arrows. Predicted Motif-1 processing sites are marked with blue dashed vertical lines. (G) Mapping of 5′ ends at the 3′ region of SACI_RS09365 (*tnpB*, IS*200*/IS*605* family) reveals two TPSs possibly associated with the generation of a ωRNA. Bar plots show normalized 5′ end read counts from +TEX/−-TEX libraries. Additionally, coverage plots of log_10_(Normalized Reads) from -TEX (dark gray) and +TEX (light yellow) show peak depletion at the positions where TPSs are identified. Predicted processing sites associated with Motif-1 and Motif-4 are indicated by blue and black dashed vertical lines, respectively.

To identify conserved sequence features potentially associated with RNA processing, we extracted 10 nucleotides upstream and downstream of each TPS and subjected the resulting sequences to motif discovery using MEME (46). This analysis revealed four significantly enriched motifs: Motif-1 (E-value = 7.5 × 10^-32^) was found 24 times in ncRNAs and 153 times in mRNAs (Fig. 2B); Motif-2 (E-value = 9 × 10^-24^) occurred 30 times in ncRNAs and 318 times in mRNAs (Fig. 2C); Motif-3 (E-value = 7.1 × 10^-17^) appeared 18 times in ncRNAs and 219 times in mRNAs (Fig. 2D); and Motif-4 (E-value = 1.2 × 10^-10^) was detected 74 times in ncRNAs and 264 times in mRNAs (Fig. 2E). Collectively, these motifs accounted for 1,100 of the 6,013 TPSs (∼18%).

Further analysis showed that Motif-1 is frequently represented in the locus of CRISPR RNA (crRNA) repeat units (98 out of 293 total Motif-1-TPSs). The transcript processing sites of manually selected crRNAs for Motif-1 revealed the accurate processing site responsible for crRNA maturation observed in the CRISPR type I-D system of *S. acidocaldarius* (Fig. 2F). Next, visualization of the 5’ ends of reads within a portion of the CRISPR array evidenced a clear depletion following TEX treatment (Fig. 2F). Moreover, analysis of -TEX coverage depicted a distinct stepwise profile, common to previously observed patterns of crRNA maturation by Cas6 (47–50).

To compare our results with previous studies, we looked specifically at the gene SACI_RS09365, which codes for a TnpB from the IS*200*/IS*605* family of insertion sequences (51). Genes of this family are known to contain an overlapping ncRNA at their 3′-end, called ωRNA, which is generated from a sequence of processing steps (41, 52, 53). Similarly to what was previously described, we also identified processing sites at the expected region. This evidence, together with an increased coverage from sRNA-seq, suggests the presence of a ωRNA (Fig. 2G). Finally, analysis of the 3′-end of the gene SACI_RS11500, which codes for a hypothetical protein and is downregulated in most conditions, presents a processing pattern and -TEX coverage that points towards the production of 3′ UTR-derived ncRNAs (54) (Fig. S1A).

Taken together with the transcriptome-wide analysis, which showed that transcripts can contain between 0 and 14 TPSs (Fig. S1B-C), these results evidence that sequence composition may act as an important determinant for recognition and/or efficiency of a subset of RNA processing and degradation events that generate 5′-monophosphorylated transcripts in *S. acidocaldarius*.

### SmAP1 and SmAP2 are essential while SmAP3 is dispensable

Previous studies have identified several RNA species as binding partners of diverse SmAPs, suggesting that these proteins participate in multiple RNA-regulated processes (20–22, 25, 27, 28). In *S. acidocaldarius*, three distinct *smAP* genes have been annotated: SACI_RS05835 (*smAP1*), SACI_RS03825 (*smAP2*), and SACI_RS03145 (*smAP3*) (Fig. S2A). The presence of 3 *smAP* genes is consistent with members of the TACK superphylum and the ASGARD group (55). To elucidate the functions of these SmAPs in *S. acidocaldarius*, we investigated: (i) the essentiality of each *smAP* gene under standard growth conditions; (ii) the transcriptome-wide determination of their RNA interaction partners via RIP-seq; and (iii) their sequence and positional binding preferences toward distinct RNAs.

To assess the essentiality of the three *S. acidocaldarius* SmAPs, we employed a targeted gene disruption strategy utilizing the organism’s endogenous type I-A and III-B CRISPR-Cas systems (56). Briefly, mutagenesis plasmids were constructed containing (i) a CRISPR array encoding a crRNA targeting the *smAP* gene of interest and (ii) homologous flanking regions designed to facilitate recombination with a donor DNA template. As negative controls, plasmids lacking the donor DNA were transformed to verify CRISPR-mediated interference activity.

Transformation of *S. acidocaldarius* with constructs targeting *smAP1* or *smAP2* resulted in sparse colony formation, regardless of whether donor DNA was included (data not shown). PCR screening of these colonies confirmed the absence of recombination events; all colonies lacked the integrated donor DNA and were classified as escape mutants or background (Fig. S2B). In contrast, transformation targeting *smAP3* yielded multiple colonies (data not shown). A PCR analysis revealed the presence of both mixed populations (wild-type and knockout alleles) and complete knockouts (Fig. S2B).

To independently validate these results, we repeated the gene disruption experiments using the established *pyrEF* counter-selection system (57). Consistent with the CRISPR-based approach, no viable transformants were obtained for *smAP1* or *smAP2*, while multiple colonies were recovered for the *smAP3* deletion. These findings collectively indicate that *smAP1* and *smAP2* are essential and non-redundant under the tested conditions, whereas *smAP3* is non-essential in *S. acidocaldarius*. This is corroborated by the essentiality classification database of the closely related organism *Saccharolobus islandicus*, where the putative homologs of SmAP1 and SmAP2 are classified as essential, whereas the putative homolog of SmAP3 is classified as non-essential (58).

### Identification of RNA Interaction Partners of SmAP1, SmAP2, and SmAP3 through Genomic Tagging and RNA Immunoprecipitation

To characterize the RNA-binding profiles of the three *S. acidocaldariu*s SmAPs, we generated genomically C-terminal His-tagged strains for each gene. All three tagged SmAP variants were successfully constructed without measurable growth defects under standard conditions (Fig. S2C).

Cultures of each strain were grown to logarithmic phase and then purified using Ni-NTA chromatography (Fig. 3A). To enrich RNA-binding partners, anti-His-tag-based RNA immunoprecipitation (RIP) was performed. PAGE analysis of the co-immunoprecipitated sample revealed substantial RNA recovery for SmAP1 and SmAP2, whereas SmAP3 yielded negligible RNA co-isolation (Fig. 3B). Therefore, we continued focusing on the interacting RNAs of SmAP1 and SmAP2 only. Total RNA from the RIP samples of SmAP1 and SmAP2 was extracted and used to construct RIP-seq libraries that were further sequenced using Illumina. The mock purification of the non-tagged wildtype was used as a background RNA control.

**Figure 3:**
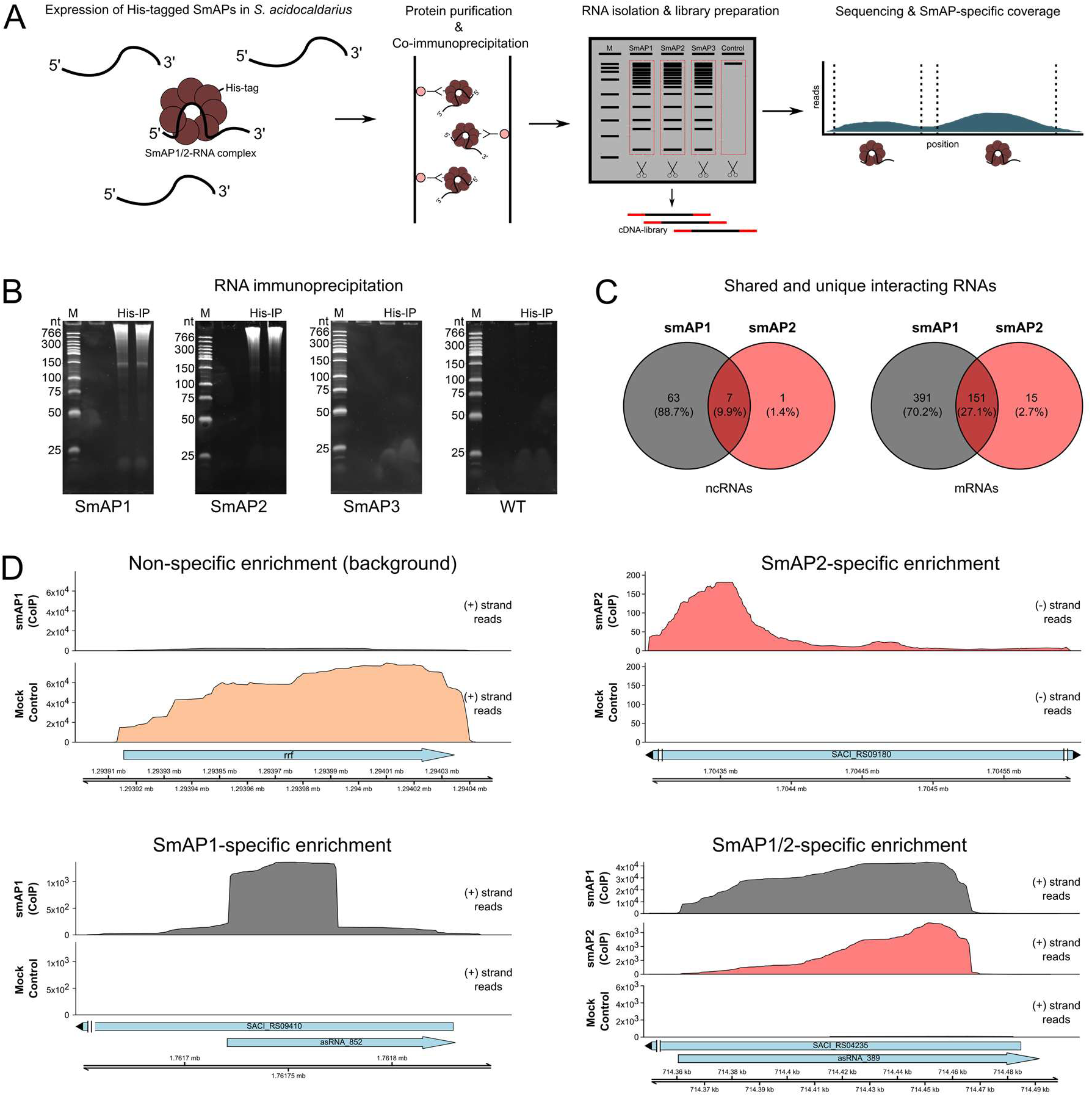
RIP-seq for the identification of SmAP interacting RNAs and their interaction sites. (A) RIP-Seq was used to identify RNAs bound by smAP1-2, combining immunoprecipitation of RNA-protein complexes with high-throughput sequencing of co-precipitated RNAs. (B) Polyacrylamide gel electrophoresis with co-purified RNA of genomically His-tagged SmAP1-3 and a mock control using *S. acidocaldarius* DSM639 MW001 (WT), evidence of abundant RNA binding for SmaP1 and SmAP2 but not for SmAP3. (C) Venn diagram showing the number of RNAs uniquely or jointly bound by SmAP1 or SmAP2. RNA binding partners were defined as significantly enriched if log_2_-fold change ≥ 1 and adjusted p-value < 0.01. (D) Coverage of RIP- seq reads for examples of coimmunoprecipitated RNAs for SmAP1 and SmAP2 individually and in combination. Reads are shown for the respective mRNA and asRNA on either the positive (+) or negative (−) strand. The mock control for background determination is depicted with the *rrf* gene (first panel). Blue arrows indicate gene orientation with left-pointing for reverse strand and right-pointing for forward strand.

In total, we identified 542 mRNAs and 70 ncRNAs, including asRNAs, as interaction partners of SmAP1. SmAP2 was associated with 166 mRNAs and 8 ncRNAs. While a subset of transcripts was bound by both proteins, most targets were unique to each SmAP, indicating specialized roles and binding specificities (Fig. 3C). Transcriptome-wide mapping of RIP-seq reads identified significant enrichment peaks, indicating direct or stable indirect RNA-protein interactions (Fig. 3D). The peaks were used to identify SmAP-bound RNAs to SmAP1 and SmAP2, either individually (e.g., asRNA_852 or SACI_RS09180 mRNA) or jointly (e.g., asRNA_389).

To explore potential binding preferences, we classified binding site positions relative to each transcript. Briefly, the length of each mRNA was normalized to 100%, and three regions were defined: the 5′ region (first 25%), the internal region (25-75%), and the 3′ region (final 25%). These regions were then used as references to analyze potential positional biases for SmAP-RNA interaction. SmAP1 displayed a predominant enrichment at the 3′ ends of mRNAs, consistent with observations in other archaeal species such as *Pyrococcus furiosus* (21). SmAP2, by contrast, showed a more balanced distribution between internal and 3′ regions, with internal sites being slightly more enriched than in SmAP1 (Fig. S2D).

For non-coding RNAs, SmAP1-associated sites were more evenly distributed between the 5′ and 3′ ends, while SmAP2 showed a marked preference for 5′ and internal regions, with fewer 3′-associated interactions (Fig. S2D). This divergence in binding location might suggest distinct functional roles in RNA metabolism or processing.

To further compare the binding characteristics of the two proteins, we analyzed the GC content of the bound regions, categorized by position. Apart from internal interaction sites for SmAP1, all categories displayed significantly reduced GC content compared to the already low (37%) genome-wide average (Fig. S2E). These findings support the hypothesis that both SmAP1 and SmAP2 bind distinct RNA populations, with differences in both target identity and positional preference. The GC content trends observed are consistent with previously reported characteristics of SmAP-bound regions in other archaeal systems, where AT-rich sequences are commonly reported (21, 25, 27, 29, 59).

### The *smAP1* gene presents evidence for a multi-layered post-transcriptional regulation

Our integrated atlas revealed a previously unannotated asRNA (asRNA_536) overlapping the locus of *smAP1* (SACI_RS05835) (Fig. 4A). Furthermore, RIP-seq analysis showed binding of the SmAP1 protein to the asRNA_536 with an interaction site mostly in its 5′ region. Inside this SmAP1 interaction site, two transcript processing sites were identified in each transcript, which could be connected to SmAP1-mediated processing of the bound asRNA (Fig. 4A). Expression analyses across different environmental and genetic backgrounds further suggest a regulatory link. The *smAP1* mRNA and asRNA_536 are generally co-regulated, evidencing a coordinated production of both. However, several NUDIX-deletion strains display an inverse pattern (Fig. 4B iii), suggesting that metabolite-sensing hydrolases might affect the equilibrium between the two transcripts. Moreover, immediately after heat stress, asRNA_536 is induced but then declines over time, whereas SmAP1 protein accumulates progressively despite relatively stable *smAP1* mRNA levels (Fig. 4C). This disconnection between protein and mRNA levels, together with an initial asRNA upregulation, points to post-transcriptional control. We propose that asRNA_536 pairs with *smAP1* mRNA, potentially leading to the decrease of translation by sterically or structurally impairing the translation machinery. As asRNA_536 levels reduce, translation is resumed, and protein levels increase. Put together, these results suggest a feedback loop in which SmAP1 binds and is translationally regulated by its asRNA.

**Figure 4:**
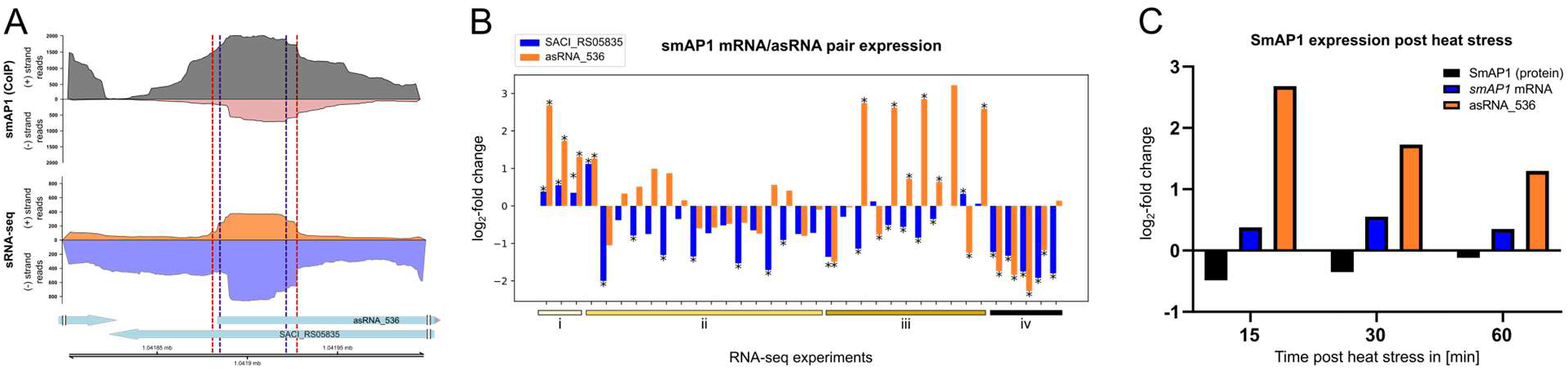
Analysis of SmAP1 (SACI_RS05835) and its asRNA (asRNA_536) expression and SmAP1 interaction site. (A) Genomic locus of smAP1 (SACI_RS05835) with the annotations for the identified asRNA_536, SmAP1 interaction site, and transcript processing sites. RNA sequencing coverage plots of the smAP1 (SACI_RS05835) locus, including reads for the SmAP1- specific enriched RNAs identified by RIP-seq (top panel) and reads identified by sRNA-seq (bottom panel). Depicted are the positive and negative strands, respectively. Transcript processing sites identified by dRNA-seq are shown in dashed lines (positive strand for blue dashed lines and negative strand for red dashed lines). (B) Differential expression profiles for the SmAP1 mRNA (SACI_RS05835) and its asRNA (asRNA_536) for various experimental conditions: i) Post heat stress (86 °C); ii) Different growth phases, temperature, and pH; iii) Knockouts of NUDIX proteins; iv) Post starvation. Significant p-values (< 0.05) obtained after DESeq2 analysis are represented as *. (C) Log_2_-fold change of SmAP1 protein abundance, *smAP1* mRNA (SACI_RS05835), and asRNA_536 levels after 15-, 30-, and 60 minutes following heat stress at 86 °C.

### Expanding the transcriptional and post-transcriptional heat stress response network in *S. acidocaldarius*

Previous studies evidenced a lack of correlation between mRNA and protein levels following heat stress in *S. acidocaldarius*, suggesting drastic effects of post-transcriptional and post-translational regulation (35). To shed light on this problem, we incorporated our expanded annotation of asRNAs to re-analyze heat stress related RNA-seq and quantitative proteomics datasets (35). To evaluate the effects of asRNAs in mRNA levels, we analyzed differentially regulated asRNA/mRNA pairs and compared the direction of their expression (Fig. 5A). At 15 minutes, most pairs cluster in the upper quadrants, showing an initial induction of asRNA expression alongside mRNA changes. By 30 minutes, there is a broader distribution, with co-regulated pairs still presenting more instances of the inverse pattern. At 60 minutes, most asRNAs remain upregulated while their cognate mRNA expression either drops further or increases, evidencing a dynamic switch from coordinated to antagonistic regulation as the stress response progresses.

**Figure 5:**
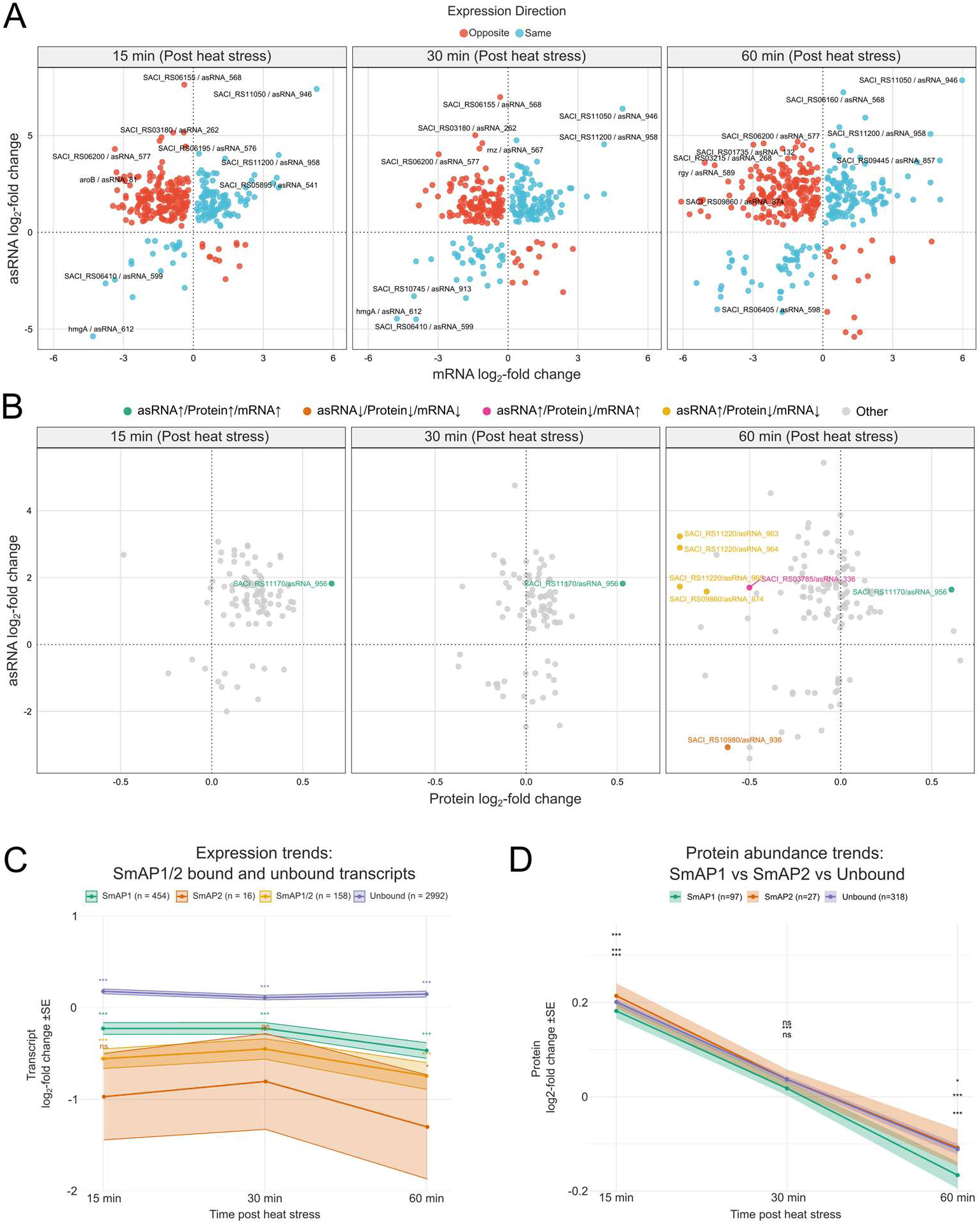
Expression analysis of mRNA/asRNA pairs with their respective protein abundance post heat stress using RNA-seq and mass spectrometry data. (A) Differential expression analysis of asRNAs in combination with their cognate mRNAs following heat stress. Depicted is the log_2_-fold change of both transcript types with either the same or opposite expression trend visualized in blue and red, respectively. (B) Differential expression analysis of asRNAs in combination with the protein levels of their cognate mRNAs following heat stress. Change in expression is represented by the log_2_-fold change, and significant expression trends are highlighted with their transcript code and the respective color for up- or downregulation. (C) Ribbon plot depicting the expression trends of SmAP1- and SmAP2-bound and unbound transcripts following heat stress. The mean expression of SmAP1/2-bound transcripts identified using RIP-seq was compared to their expression levels observed after heat stress. Change in expression is represented by the log_2_-fold change. (D) Ribbon plot depicting the mean protein abundance after translation of SmAP1- and SmAP2-bound and unbound mRNAs following heat stress. Change in expression is represented by the log_2_-fold change after comparison of the post and pre heat stress conditions. Significant p-values are represented by * p < 0.05, *** p < 0.005, ns: not significant.

Next, to examine how asRNAs coordinate with protein-level changes after heat stress, we plotted asRNA log₂-fold changes (y-axis) against protein log₂-fold changes (x-axis) at 15-,30- and 60-minutes post-stress (Fig. 5B). At 15 minutes, nearly all points cluster around the origin, with only SACI_RS11170/asRNA_956 appearing as an outlier, pointing to increased asRNA, mRNA, and protein levels. By 30 minutes, most pairs shift towards the origin, while SACI_RS11170/asRNA_956 maintains its coordinated upregulation. At 60 minutes, however, a more diverse scenario is visualized; several genes reflect concurrent downregulation (Fig. 5B), while other outliers, such as SACI_RS03785/asRNA_336, present a rise in mRNA and asRNA levels but decreased protein levels, suggesting an antisense-mediated translational repression. Finally, cases where the asRNA is upregulated, but the protein and mRNA are less abundant (e.g., SACI_RS11220/asRNA_963/964), suggest that asRNAs may accelerate mRNA decay, indirectly dampening protein synthesis. These results reveal that, whereas the earliest response is largely uniform, as it progresses, a more diverse scenario consistent with antisense-driven post-transcriptional mechanisms can be suggested.

Transcriptome profiling reveals that SmAP2-bound RNAs undergo the strongest and most persistent downregulation after heat stress, with SmAP1-only targets and co-bound (SmAP1/2) transcripts showing intermediate repression at early and late timepoints (Fig. 5C). On the other hand, unbound transcripts tend to show upregulation at all three timepoints. At the protein level, however, a biphasic protein response that is largely independent of SmAP-binding status is detected (Fig. 5D). These divergent mRNA and protein trajectories suggest that SmAP association might lead to sustained post-transcriptional repression.

### Creation of the *S. acidocaldarius* transcriptomic atlas

To provide an easy-to-access resource for the community, a web-based atlas, built upon our manually curated genome annotation of *S. acidocaldarius* that incorporates multiple publicly available and previously unpublished transcriptomic datasets, was created (Fig. 6). Users can compare differential RNA-seq results across multiple conditions and genotypes. Visualizations are offered in both tabular format and as fold-change-specific graphs, enabling intuitive and scalable exploration of gene expression dynamics. To explore the impacts of RNA-mediated regulation, we implemented an asRNA/mRNA pair explorer. This tool allows users to assess expression correlations between asRNAs and their cognate genes across datasets, with the potential to identify condition-specific regulatory relationships relevant to stress adaptation or genetic perturbation. The atlas also includes dedicated modules to explore TPS motifs and SmAP interaction sites found through RIP-seq. These features allow for the visualization and localization of enriched RNA-binding regions and processing signatures directly within the genome browser interface. An integrated IGV-based genome viewer enables interactive browsing of annotated features and transcriptomic tracks, easing the inspection of specific loci. To further support functional interpretation, we incorporated a Clusters of Orthologous Groups (arCOG) assignment module that matches *S. acidocaldarius* genes with their respective arCOG class. By combining diverse transcriptomic data into a unified and user-friendly platform, this atlas is a valuable community resource for exploring post-transcriptional regulation in *S. acidocaldariu*s. It also sets up a framework that can be downloaded and locally customized for other archaeal systems for comparative analysis. The atlas is available at (https://vicentebr.github.io/Sulfolobus_atlas/).

**Figure 6:**
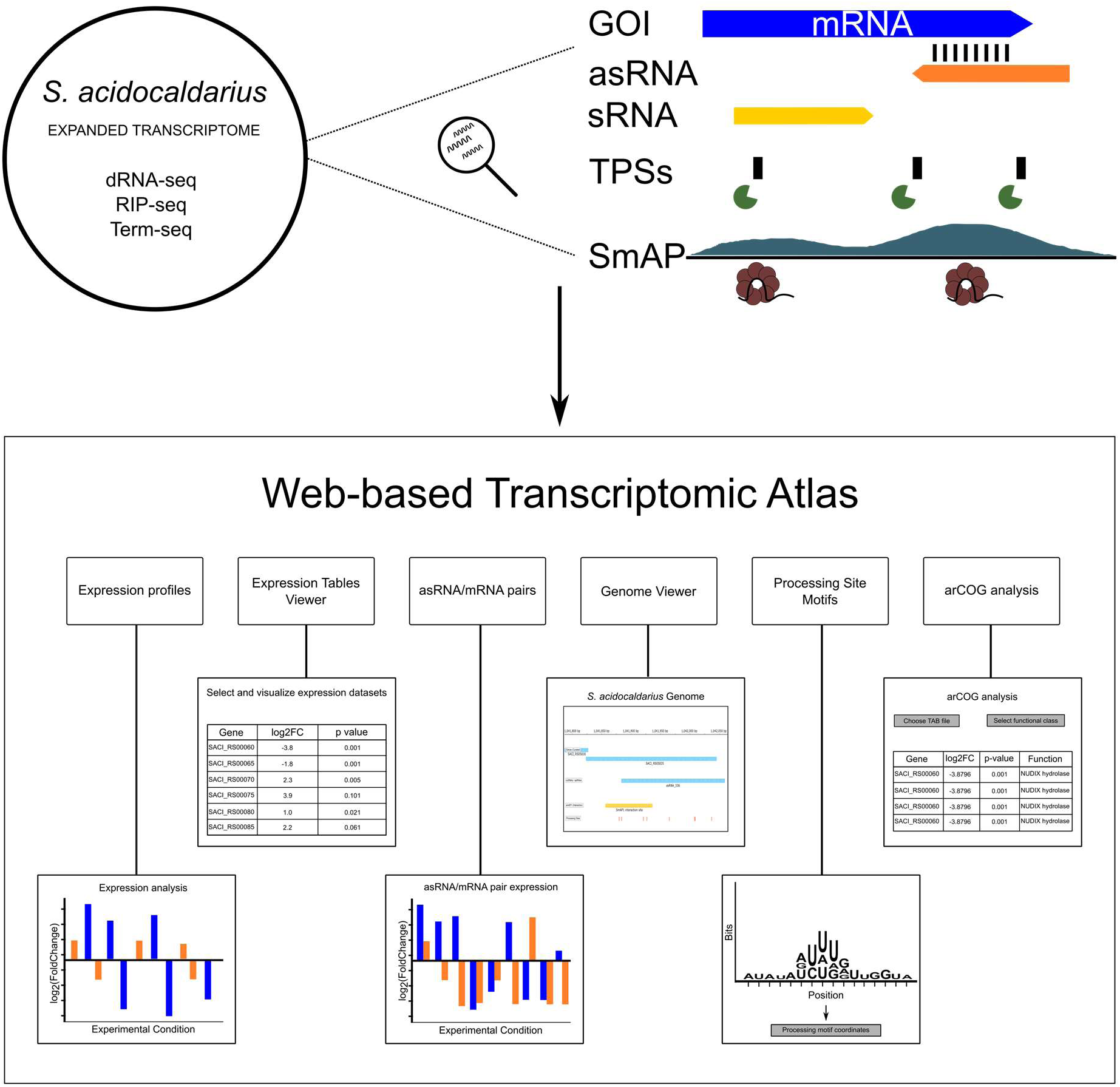
Schematic overview of the transcriptomic atlas based on the combiation of various RNA-sequencing strategies for exploring transcriptomic data over different conditions. Depiction of the framework that was used to construct a comprehensive transcriptomic atlas based on both public and newly generated RNA-seq datasets across multiple experimental conditions. The integrated data enables the identification of novel asRNAs, sRNAs, TPSs, and RNA-binding partners of SmAP1 and SmAP2. Combined analysis of these datasets supports the discovery of putative post-transcriptional regulatory events associated with specific genes of interest (GOIs). The data generated is then combined and made accessible in a web interface that includes six key modules: i) Expression Profiles: generates plots of the expression profile of a specific gene across multiple experimental conditions. ii) Expression Tables Viewer: allows users to visualize and select which datasets will be plotted in the Expression Profile module. iii) asRNA/mRNA Pair Explorer: allows the visualization of the expression profiles of mRNAs and their corresponding asRNAs. Correlation information is also provided. iv) Genome Viewer: displays *S. acidocaldarius* DSM 639 curated annotation tracks for genes, mobile elements, ncRNAs/asRNAs, SmAP1 and SmAP2 binding sites, and transcript processing motifs. v) Processing Site Motifs: allows visualization of enriched sequence motifs surrounding predicted transcript processing sites, their sequences, and position in the genome. vi) arCOG analysis: links gene expression datasets (e.g., DESeq2 outputs) to arCOGs for functional characterization and pathway analysis.

## DISCUSSION

The transcriptome-wide identification of RNA interaction partners for SmAP1 and SmAP2, together with the global mapping of TPSs and asRNAs, sheds light on the post-transcriptional regulatory features of *S. acidocaldarius*. Our updated transcript annotation expands the non-coding RNA repertoire of *S. acidocaldarius*, uncovering 990 asRNAs and 160 ncRNAs. This increase aligns with similar findings in other archaea, such as *Haloferax volcanii*, *H. salinarum*, and *Methanosarcina mazei*, where high-throughput sequencing technologies have revealed pervasive transcription and extensive antisense regulation (9, 39, 60). The distribution of asRNAs suggests a potential role in modulating transcript stability or translation, consistent with known mechanisms in archaea and bacteria (13–15, 24, 39, 61, 62). The identification of a conserved TATA-box motif upstream of asRNAs and ncRNAs reinforces that these transcripts are most likely regulated and not products of spurious transcription (63, 64).

Expression correlation analyses between asRNAs and their cognate mRNAs reveal both positively and negatively correlated pairs. Negatively correlated examples, such as asRNA_480 and its cognate ABC transporter gene SACI_RS05085, point to a classic asRNA-mediated repression, which has been reported for transporter systems in hyperthermophilic archaea (65). In contrast, positively correlated pairs, such as asRNA_559 and the sulfolobicin gene SACI_RS06075, may suggest co-regulation or effects on RNA stability (66, 67).

Our transcriptome-wide analysis of TPSs uncovered over 6000 processing events, with motif analysis revealing four conserved sequence elements enriched at processing sites. These motifs exhibit distinct positional preferences, especially Motif-1, which is biased toward 5′ regions in mRNAs and internal regions in ncRNAs. The presence in crRNA repeat units, potentially within the recognition site for the endoribonuclease Cas6 of the CRISPR type I-D system, reinforces its relevance for RNA maturation (49, 68). Similar processing signatures are suggested to influence transcript stability and maturation in other archaeal species, further supporting the hypothesis that TPSs contribute significantly to post-transcriptional regulation in Archaea (41, 66).

SmAPs represent a central hub of post-transcriptional control. Knockout attempts on *S. acidocaldarius* revealed that SmAP1 and SmAP2 are essential, while SmAP3 is dispensable under standard growth conditions. RIP-seq mapping of SmAP-RNA interactions showed that SmAP1 binds a broader set of RNAs, at their 3′ ends, consistent with its hypothesized role in RNA stabilization or decay. It has been shown that SmAP homologs in *Sa. solfataricus* performs physical interactions with the exosome protein complex (25, 27). The binding of SmAP1 to 3′ ends could be linked to exosome recruitment, potentially leading to 3′-5′-directed RNA degradation (27). SmAP2, although interacting with fewer RNAs, shows a higher internal binding enrichment. These findings are consistent with earlier structural and biochemical studies of SmAPs in *Sa. solfataricus* and *P. abyssi*, which demonstrated differential binding affinities and functions among SmAP paralogs (21, 25, 59). The RIP-seq analysis of SmAP1-specific transcripts also suggested asRNA-related mechanisms corresponding to heat stress. The asRNA_536 of the locus of SmAP1, which is also bound by SmAP1 itself, seems to be involved in heat stress response. We propose that SmAP1 is regulated by its asRNA in a mRNA titration manner, with the possibility to inhibit translation. The overall inhibition of SmAP1 mRNA translation in the early phase of heat stress could be part of a major stress response regulation system.

Earlier observations that transcript- and protein-level responses to heat stress in *S. acidocaldarius* exhibit limited overlap, as shown by a gene-by-gene correlation analysis, suggested that a time-delayed proteome response in the same direction as the transcriptome is not the most likely scenario, and that post-transcriptional and post-translational regulation might be prevalent (35). In this sense, our expanded asRNA annotation, together with the SmAP1 and SmAP2 binding profiles, would be valuable additions to shed light on this problem. Indeed, we observed that positively or negatively correlated asRNA/mRNA pairs are detected after heat stress. Moreover, when we map these dynamics into proteomic changes, we uncovered potential candidates for antisense-mediated translational repression and antisense-driven mRNA decay that cannot be predicted without proper annotation of asRNAs. Additionally, transcripts bound by SmAP1 or SmAP2 display a sustained downregulation when compared to unbound transcripts. Protein levels, on the other hand, do not seem to be affected in a SmAP1- or SmAP2-dependent manner, as the trends of both bound and unbound transcripts follow a similar pattern. Together, these findings expand and help to explain the observed disconnect between mRNA- and protein-levels and establish asRNAs and SmAPs as potential key regulators during the *S. acidocaldarius* heat shock response.

While our evidence is primarily correlative, the integrated atlas and the described loci provide starting points for mechanistic studies. Targeted RNA duplex disruption, ribosome profiling, RNase mapping, and conditional SmAP depletion (e.g., gene knockdown) will be key to demonstrating causality. Altogether, this work establishes asRNA and SmAPs as important condition-dependent regulators of gene expression in *S. acidocaldarius* and provides a community resource to facilitate data exploration and hypothesis generation.

## MATERIALS AND METHODS

### Strains, Plasmids, and Primers

All strains, plasmids, and oligonucleotide sequences used in this study are described in Table S1. This work used *S. acidocaldariu*s DSM639 MW001 (MW001 for short), an uracil-auxotroph strain. Cultures were grown aerobically at 120 rpm and 75 °C in Brock medium, pH 3.5. The medium was supplied with 0.1% (w/v) NZ-Amine and 0.2% (w/v) dextrin, and 10 µg/ml uracil. *E. coli* strains were grown aerobically at 180 rpm and 37 °C in LB medium (0.5% (w/v) yeast extract, 1% (w/v) tryptone, 1% (w/v) NaCl). For solid medium, LB medium was mixed with 1.5% (w/v) agar-agar and supplied with the respective antibiotic (0.001% (v/v)). Cell growth was achieved by monitoring the optical density of the cultures at 600 nm.

### General analysis for publicly available RNA-seq data

Previously published transcriptome datasets (Table S3) were downloaded from the ENA database and reanalyzed. Briefly, raw reads were quality and adapter trimmed using Cutadapt (v2.8) and quality checked with FASTQC (v0.11.9). Processed reads (≥18 nt) were mapped to the reference genome of either *S. acidocaldarius* using Hisat2 (v2.2.1). After strand-specific screening, featureCounts was used to count gene hits. Statistical and enrichment analyses were performed with DESeq2 (v1.36.0). Genes with log2-fold change ≥ 1 or ≤ −1, with p-value < 0.05 were considered differentially expressed. Normalized bedgraph genome coverage files were generated using bedtools. The Integrative Genomics Viewer (IGV, v2.13.2) was used to inspect and visualize candidate sequences.

### Expanded non-coding RNA and smORF annotation in *S. acidocaldarius*

Previously published dRNA-seq (PRJEB48624) (3) and Term-Seq (33) datasets for *S. acidocaldarius* were reanalyzed to refine gene annotation. Transcription start sites (TSSs) were identified using TSSAR (Transcription Start Site Annotation Regime) Web Service (v1457945232) (43). Following initial annotation, TSSs categorized as orphan (oTSS) or antisense (asTSS) were filtered based on the following criteria: score ≥ 600; positional difference ≥ 10; p-value ≤ 0.05.

To determine the boundaries of newly identified transcripts, annotated transcription termination sites (TTSs) were searched within a 400 bp window downstream of each oTSS and asTSS. Next, the expression of the transcript candidates was evaluated over multiple datasets (File S2). The coding potential of each transcript was assessed by identifying potential start and stop codons within the annotated regions. Transcripts with validated expressions under at least one analyzed condition were incorporated into the final annotation. The updated *S. acidocaldarius* transcriptome annotation is provided in File S1.

### Identification of putative transcript processing sites (TPS) in *S. acidocaldarius*

Identification of putative transcript processing sites was performed similarly to REF. Briefly, TSSAR was used to identify TPS positions by inverting the TEX+ and TEX− input files. Thus, the identified positions are likely 5′-p positions that were depleted after TEX treatment. TSSAR parameters were p-value (p) of p < 10−9 with a minimum of 10 reads and “TSS” grouping of at least 5 nt.) Following initial detection, TPSs were filtered based on the following criteria: score ≥ 600; positional difference ≥ 10; p-value ≤ 0.05.

To identify motifs potentially associated with processing, the position of each TPS was extended 10 nts upstream and downstream, and the sequence extracted with bedtools (69) Identification of motifs was performed using Meme (-mod zoops -minw 4 -maxw 40 -objfun classic -markov_order 0) and Glam2 (46).

### Construction of chromosomally 6xHis-tagged *smAP1*, *smAP2*, *smAP3* strains in *S. acidocaldarius*

Genomic His tagging of the *smAP* genes was performed using the pop-in/pop-out method based on the single crossover recombination steps with the pSVA406 plasmid (57). *S. acidocaldarius* MW001 cells were transformed with the respective plasmids (Table S1). Transformants were grown to the logarithmic phase in 50 ml Brock media supplemented with 0.1 % (w/v) tryptone and 0.2 % (w/v) sucrose. 10 μl of the cell cultures were diluted in 100 μl ddH_2_O and plated on second selection plates bearing 5-FOA and uracil. The 5-FOA induced the loss of the plasmid, yielding a 50 % chance of the aimed mutation. The plates were incubated for 5 days, and the obtained colonies were screened for correct mutations by colony PCR and sequencing.

### Construction of a marker-less smAP3 gene deletion strain in *S. acidocaldarius*

The marker-less *smAP3* gene deletion was performed using the CRISPR-Cas type I-A/III-B interference system (56). The endogenous CRISPR-Cas type I-A/III-B system of *Sulfolobus acidocaldarius* allows a plasmid-based gene deletion via recombination by providing the target gene flanking donor DNA and a gene-targeting crRNA. Briefly, the target gene deletion is acquired by the preceding recombination induced by the donor DNA provided. Only cells with prior recombination can survive, while cells without recombination are self-targeted by the plasmid-based CRISPR system, causing DNA cuts with subsequent DNA degradation and cell death. For the cleavage of the *smAP3* target DNA by the endogenous CRISPR-Cas system, a 38 nt spacer and a TCT PAM were chosen. The spacer sequence was cloned into the pSVAxylFX-CRISPR base vector, and expression of the crRNAs was achieved by supplementing the culture plates with 0.3 % (w/v) D-xylose using the pentose-inducible *saci_1938* promoter. The deletion of *smAP3* was verified by colony PCR with target-flanking primers and sequencing (Whole Genome Sequencing, Novogene). The same approach was used for the *smAP1* and *smAP2* genes, but this did not result in gene deletion (Fig. S2B). Gene disruption was also attempted using the *pyrEF*-exchange method. The *pyrEF* gene cassette of *Sa. solfataricus*, amplified using primer pairs with overhangs that are homologous to the target gene, is electroporated and subsequently integrated into the *S. acidocaldarius* genome via homologous recombination. Successful genomic manipulation results in clones growing in a medium that does not contain uracil. As with the previous approach, only the *smAP3* gene could be successfully disrupted.

### smAP1-3-RNA coimmunoprecipitation

Two subsequent purification steps were conducted to isolate the interacting RNAs of the SmAP proteins: i) Ni-NTA purification and ii) immunoprecipitation. For this, triplicates of 2-2.5 g *S. acidocaldarius* smAP1/2/3-CHis or MW001 (control) cells were resuspended in purification buffer (100 mM HEPES pH 8 for SmAP1, 100 mM MOPS pH 6.5 for SmAP2/3, 100 mM NaCl, 10 % (v/v) glycerol, 10 mM imidazole, 5 ml/g cells) and lysed by French Pressure (three times at 25,000 psi). After centrifugation of the cell lysate (30,000 × g, 20 min, 4°C), the filtered supernatant (Millex ® syringe filter, pore size 20 μm) was cycled onto a 1 ml HisTrap HP Ni-NTA column for 45 min at 4°C. The loaded column was afterwards washed with a wash buffer (purification buffer containing 25 mM imidazole) using an FPLC system. The SmAP proteins were eluted from the column by a 500 mM imidazole gradient using an elution buffer (purification buffer containing 500 mM imidazole). The four fractions that displayed the highest purity and protein amount in the SDS-PAGE analysis were pooled. 30 μg of THE^TM^ His-tag IgG antibody (from mouse) was added, the mixture was divided into two 2 ml tubes and incubated for 1 h at 4°C on a rotary shaker. 100 μl of washed Protein G Dynabeads^TM^ were added to each of the tubes, and the mixture was incubated for 20 min at 25°C. Afterwards, the magnetic beads, which were bound by the smAP-CHis-anti-His-Ab complex, were separated using a magnetic rack. The beads were washed three times with 1 ml His-IP wash buffer (purification buffer containing 250 mM imidazole). Afterwards, the beads of each tube were resuspended in a 1 ml bead wash buffer and pooled into one tube. The beads were again separated using the magnetic stand, the supernatant was removed, and the smAP-bound RNAs were eluted with 100 μl 8 M urea by incubation for 15 min at RT (shaking at 1400 rpm in a thermo mixer). All steps were performed on ice or at 4°C to avoid degradation of the RNA, unless otherwise stated. To elute proteins, a second elution was performed with 64 μl SDS loading buffer (75 mM Tris-HCl pH 6.8, 0.6 % (w/v) SDS, 0.001 % (w/v) bromophenol blue, 15 % (v/v) glycerol, 1 M β-mercaptoethanol) by incubation for 5 min at 95°C (shaking at 1400 rpm in a thermo mixer). The smAP interacting RNAs of the urea eluate were afterwards separated by denaturing PAGE and purified via gel extraction. The SDS eluate was analyzed by SDS-PAGE. The co-immunoprecipitated RNAs of SmAP proteins (RIP-Seq) were used to prepare cDNA libraries for Illumina sequencing. The library preparation was performed using the NEBNext ® Multiplex Small RNA Library Prep Set for Illumina according to the instructions of the manufacturer. The libraries were sequenced on a HiSeq2500 in single-end mode with 100 nt read lengths.

### SmAP1-2 RIP-seq data analysis

The obtained libraries were analyzed using the Ripper pipeline with a few modifications (70) Briefly, low-quality ends and adapter sequences were removed from raw reads with Cutadapt (71). Next, reads were aligned to the reference genome of *S. acidocaldarius* (NCBI_Assembly: GCF_000012285.1) using HISAT2 (-k 1000, -no spliced-alignment, -no-softclip) (72). Alignment files were converted to BAM with samtools (73) and normalized with MMR (74). A per-base coverage of the genome was generated with bedtools (69) and used as an input for a coordinate-wise log_2_-fold change (coIP vs control). Regions with at least 10 consecutive nucleotides that had a log_2_-fold change ≥ 1 were annotated as potential interaction sites. Finally, the enriched regions were tested for significance (log_2_-fold change ≥ 1 and padj < 0.01) using DESeq2 (38). To assess whether regions identified by RIP-seq presented distinct GC content compared to the genomic average, we calculated the GC content of each enriched RIP-seq peak. Next, for background comparison, the genome-wide average GC content was calculated using a sliding window (50 nt with a 25 nt step) across the *S. acidocaldarius* DSM 639 genome. A non-parametric Mann-Whitney U test was performed to determine whether the GC content distribution of RIP-seq peaks was significantly different from the background (p-value < 0.05).

### *S. acidocaldarius* Web-Based Atlas for Exploring Post-Transcriptional Regulation

The available web atlas was developed by integrating standard web technologies (HTML and CSS) and Gemini (75), an AI assistant, to assist with code snippet generation and debugging. The interactive functionalities were developed using JavaScript with the following modules: i) ApexChart for plotting dynamic and interactive bar charts with gene expression (76); ii) PapaParse for client-side parsing of all text files containing the generated data (77); iii) IGV.js for interactive genome browsing integration (78).

## Data availability

All data supporting the web application, including genomic annotations, pre-computed expression profiles, and sequence motifs, are stored and made accessible via a GitHub repository https://github.com/VicenteBR/Sulfolobus_atlas and at Zenodo (10.5281/zenodo.16312244). The raw RIP-seq data are deposited at the European Nucleotide Archive (ENA) under the accession number PRJEB94137.

## ACKNOWLEDGMENTS

We are grateful to Dr. Ankan Banerjee for his feedback and criticism. We also thank the Hessisches Kompetenzzentrum für Hochleistungrechnen (HKHLR) team for the maintenance and support during the use of the MaRC3a Cluster Marburg.

Support was provided by GRK 2937 (project number 505997786) funded by the German Research Foundation (DFG) and a Volkswagen Momentum grant to LR.

## AUTHOR CONTRIBUTION

M.B., Methodology, Formal Analysis, Investigation, Data Curation, Writing - Original Draft, Writing - Review and Editing. M.D., Methodology, Validation, Investigation. L.R., Conceptualization, Resources, Supervision, Project Administration, Funding Acquisition, Writing - Review & Editing. J.V.G-F., Conceptualization, Supervision, Formal Analysis, Data Curation, Visualization, Writing - Original Draft, Writing - Review & Editing, Software.

## FIGURES

**Figure S1:**
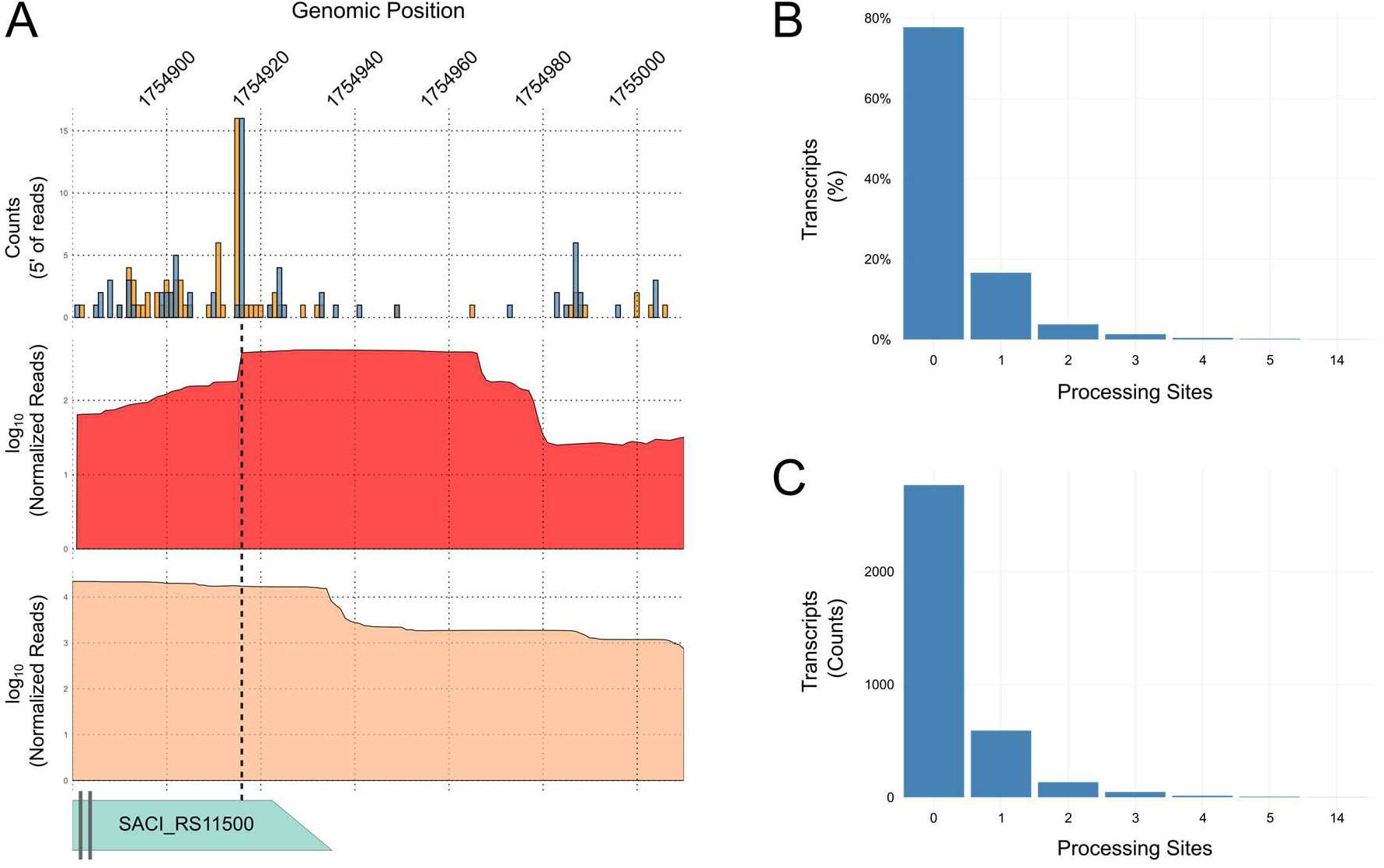
Analysis of the transcript processing site motifs identified with dRNA-seq. A) Mapping of 5′ ends at the 3′ end of the gene SACI_RS11500 reveals one TPS possibly associated with the production of 3′-UTR derived ncRNAs. Bar plots show normalized 5′ end read counts from +TEX/−TEX libraries. Additionally, coverage plots of log_10_(Normalized Reads) from -TEX (red) and +TEX (salmon) show peak depletion at the positions where TPSs are identified. B) Distribution and accumulation of TPSs per transcript, represented as %. C) Distribution and accumulation of TPSs per transcript, represented as raw counts.

**Figure S2:**
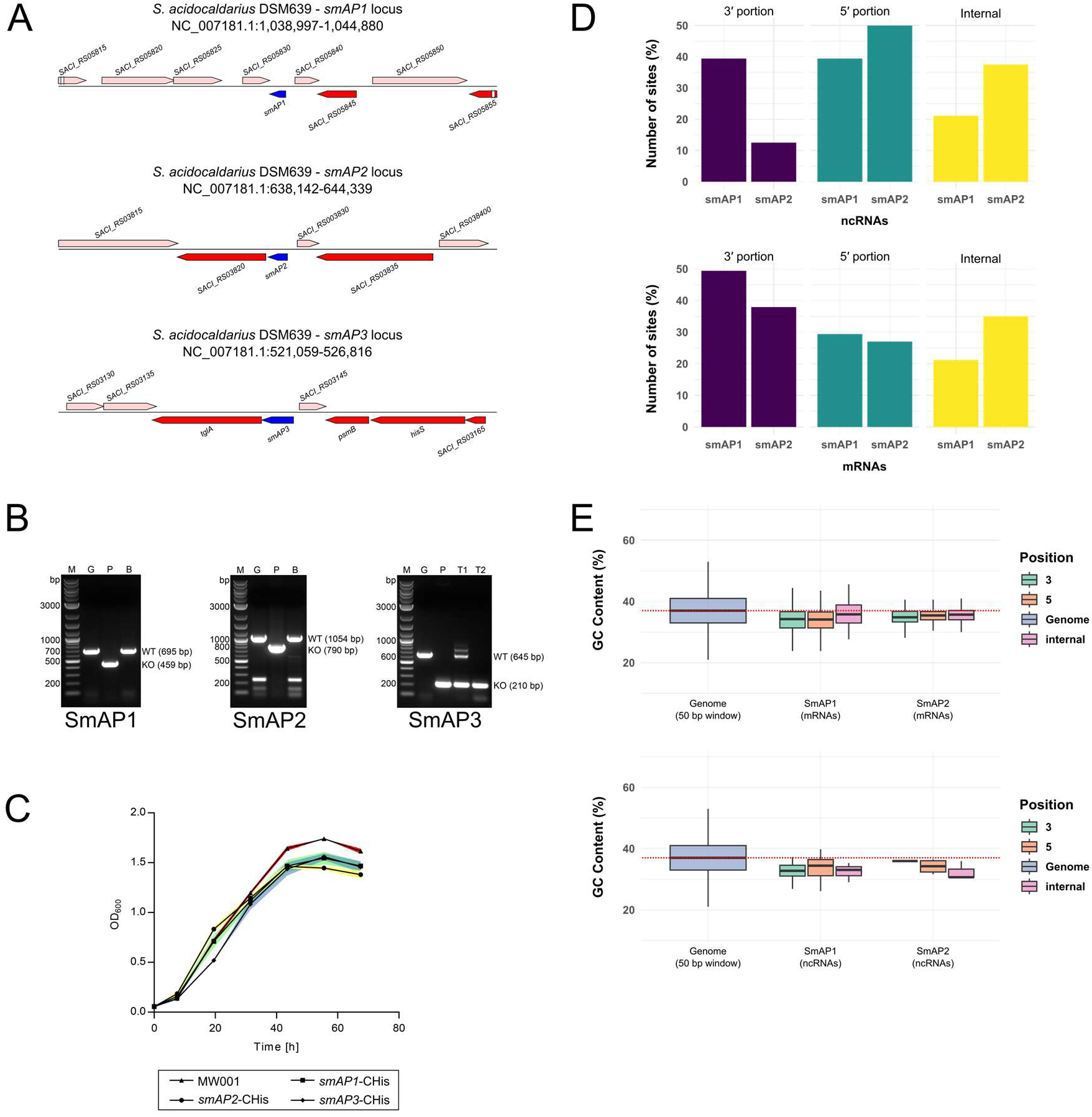
Loci organization, knockout attempts and binding properties of SmAPs. A) The *smAP1-3* loci. Blue colored arrows: the *smAP* genes. Salmon colored arrows: forward strand genes. Red colored arrows: reverse strand genes. B) Colony PCRs of the *smAP* loci show bands representing the wild-type locus (WT) or the *smAP* deleted locus (KO). Genomic DNA (marked G) of *S. acidocaldarius* and the pCRISPR plasmids (marked P) were used as negative and positive controls, respectively. Representative background colonies (marked B) of the *smAP1* and *smAP2* deletion attempts display WT bands. Two types of transformants (T1/T2) were obtained for the *smAP3* deletion attempt: T1 shows the WT and KO bands as well as additional upper bands, whereas T2 displays a single band for the *smAP3* deleted locus. C) The growth behavior of genomically His-tagged SmAP strains was compared to the *S. acidocaldarius* DSM 639 MW001 reference strain. Error bars (color-filled area) demonstrate the standard deviation of three technical replicates. D) positional biases of SmAP1-2 binding. GC content and localization on the transcript of SmAP1-2 on mRNAs (E) and ncRNAs (F).

## Supplementary Files – Legends

File S1 – *Sulfolobus acidocaldarius* DSM639 curated annotation. GFF file containing the most up-to-date RefSeq annotation and the transcripts identified in this study.

File S2 – Differential expression analysis using DESeq2 of all analyzed conditions

File S3 – Transcript processing sites positions and sequences in their vicinity (+10 and −10 nucleotides from the TPS position)

## Supplementary Tables – Legends

Table S1 – List and specification of each plasmid, strain, and oligos used in this work

Table S2 – Table of all mRNAs that have at least one associated asRNA

Table S3 – All sources of publicly available RNA-seq data, associated publication (if available), and conditions.

